# Associations Between Frailty and Cognitive Outcomes Across the Lifespan of Mice

**DOI:** 10.1101/2025.10.22.683969

**Authors:** Sydney Bartman, Lauren Gaspar, Alice Kane, Emily Potts, David A. Sinclair, Giuseppe Coppotelli, Jaime M. Ross

**Affiliations:** George and Anne Ryan Institute for Neuroscience, University of Rhode Island, Kingston, RI USA; Department of Biomedical and Pharmaceutical Sciences, College of Pharmacy, University of Rhode Island, Kingston, RI USA; Department of Genetics, Blavatnik Institute, Paul F. Glenn Center for the Biology of Aging, Harvard Medical School, Boston, MA, USA

**Keywords:** aging, frailty, behavior, sex differences, rodents

## Abstract

Advancements in monitoring biological and brain aging with precise measures of health and longevity have the potential to accelerate research on pharmacological, genetic, and aging-related interventions. Over the past decade, frailty index has been used as an assessment tool for rodents, evaluating more than 30 non-invasive parameters that are strongly associated with chronological age, correlated with mortality, and sensitive to lifespan-altering interventions. However, whether aging phenotypes captured by the frailty index reflect brain aging remains unclear. In this study, we examined the relationship between frailty index and cognitive ability in young (3–4 months), middle-aged (12 months), and old (24 months) male and female C57BL/6J mice using a battery of behavioral and locomotor assays to determine whether frailty index scores can predict performance in tasks evaluating behavioral and cognitive function. Among the behavioral assays tested, frailty index scores had good correlation with the percentage of time spent in the center of the open-field apparatus, the duration spent in the open arms of the elevated plus maze, and the time spent in the target hole of the Barnes maze. These findings indicate that the frailty index not only reflects general physiological aging but may also serve as a reliable predictor of age-related cognitive decline in mice, providing a valuable tool for studies of interventions targeting brain aging.

## Introduction

As the average age of the western world population is increasing, countries are predicting significant demographic changes over the next two-to-three decades, with the older population, defined as persons older than 65 years, expected to grow to 21-30% of the population [1,2]. Although these are projections, the increasing proportions of older individuals constitute a significant challenge [3]. Of all the aging disorders, aging of the human brain is a cause of cognitive decline and the major risk factor for neurodegenerative diseases [4,5]. Brain aging disorders represent one of the major causes of disability and dependency among the aged population and have a tremendous impact on patients, families, caregivers, and society with physical, emotional, social, and economic ramifications [6,7]. Alzheimer’s Disease (AD) and Alzheimer’s Disease Related Dementia (ADRD) comprise the most common forms of dementia and have emerged as the greatest health threats, with nearly 10 million new cases each year, currently totaling 55 million people worldwide [8].

In recent years there has been a dramatic advance in monitoring biological age, versus chronological age, by using methylation clocks [9–13]. This method, although very precise, is invasive, expensive, and time-consuming [14]. Thus, other non-invasive methodologies have been developed to monitor biological aging. One of these is frailty index (FI), a non-invasive, reliable metric for assessing the phenotypic aspects of aging using 31 different clinical-based assessments that has been applied to both humans [15] and reverse-translated to mice [16]. Frailty is known as an increase in vulnerability to negative health outcomes, and in humans, individuals possessing higher frailty indexes will have a higher demand in healthcare needs and an increased risk of mortality [18]. FI has been shown to correlate with chronological age, mortality, several age-related diseases, and has been used to detect responsiveness to health interventions [19,20]. In mice, FI has been shown to be a very powerful tool not only to determine the biological age but also for predicting lifespan [17].

Though the frailty index is a reliable tool to assess physical aging in mice, it doesn’t provide information about how the brain ages. Indeed, to monitor brain aging in mice, several behavioral tests are commonly applied that mainly measure memory deficits, such as Barnes maze, Morris water maze, and novel object recognition [21–24]. These tests are laborious and require specialized equipment and personnel; thus, it would be useful to understand how well the FI scoring system can predict cognitive decline in mice [25]. To address this question, we assessed the frailty index in healthy young, middle-aged, and old C57BL/6J male and female mice, alongside several behavioral and locomotor assays including the open-field, Y-maze, elevated plus maze, and Barnes maze, to test, for the first time, whether the FI score correlates with behavioral outcomes, thereby providing a potential measure of brain aging.

## Materials and Methods

### Animals

Both female and male C57BL/6J mice (N=120 mice in total, N=20 per sex per age group: young, middle, old) were obtained from the National Institute on Aging (NIA) Aged Rodent Colonies (Charles River Laboratories, Kingston, NY or Raleigh, NC, USA). All mice were acclimated for at least 1 month prior to testing. Testing began for the three age groups at 3, 12, 24 months, respectively, in the following order: open-field, elevated-plus maze (EPM), Y-maze, Barnes maze, and frailty index (FI). All mice received a standard diet (5053 - PicoLab® Rodent Diet 20, LabDiet, St. Louis, MO, USA) and water *ad libitum*. Mice were group-housed (3-4 mice) by sex in ventilated cages with access to a small house and nesting materials and kept on a 12:12 light:dark cycle at 21-22 °C with 45–50% humidity. Adequate measures were taken to minimize pain and discomfort. Experiments were conducted in accordance with the ethical standards and according to the Declaration of Helsinki and national and international guidelines and has been approved by the authors’ institutional review board.

### Behavioral Experiments

Behavioral tests were performed in a neutral, quiet environment between 9:00 and 17:00 (light phase) by the same researchers. Testing of the mice from the different age cohorts and sexes was staggered appropriately. Mice were acclimated in their home cage for 1 h in the testing room at 21-22 °C with 45–50% humidity prior to testing. A non-transparent plastic container, cleaned with 70% ethanol after each use, was used to transport the testing mouse to and from the apparatuses. All apparatuses were cleaned extensively with 70% ethanol after each test in order to remove olfactory cues. Tests were performed in the following order: open-field, elevated-plus maze (EPM), Y-maze, Barnes maze, and frailty index (FI). All tests were recorded and analyzed using the same software (TopScanLite v. 2, CleverSys Inc. Reston, VA, USA).

### Open-Field Assay

A chamber (48.5 cm x 48.5 cm) with darkened walls and open top was used to assess exploratory behavior and spontaneous locomotion (CleverSys Inc., Reston, VA, USA). During this test, mice were placed in the chambers and allowed to explore for 1 h. The open-field apparatus used consisted of four chambers thus allowing four mice to be tested at the same time. Total distance traveled and duration in center versus the periphery were recorded.

### Elevated-Plus Maze

The elevated plus maze (EPM) consists of a cross-shaped platform with four arms, two with walls and two lacking walls, that is elevated from the floor. The EPM used had arms measuring 7 cm in width, 30 cm in length, with a wall height of 15 cm, and was 59 cm in height from the floor (CleverSys Inc., Reston, VA, USA) [26]. Each mouse was placed in the center of the EPM and allowed to explore for 5 min. The duration spent and the number of bouts in the open arms were measured.

### Y-Maze

A dry maze comprised of three symmetrical arms that resemble a Y-shape, with the ability to close off access to one of the arms was used (CleverSys Inc., Reston, VA, USA). During the first trial, each mouse was allowed 3 min to explore the maze with one of three arms closed. In between trials, mice rested in their home cages for 15 min. During the second trial, the previously closed arm was open, and each mouse was allowed an additional 3 min to explore all three arms. The duration spent and the number of bouts in the novel arm were measured.

### Barnes Maze

A dry maze consisting of an elevated (82 cm) white circular platform (90 cm in diameter) with 20 x 5 cm diameter holes evenly spaced along the circumference divided into four quadrants was used. One of these holes, the target (T), held an escape box underneath the maze in which a mouse could fit. Four visual cues of different colors and patterns were placed on the walls surrounding the maze and bright lights were used as a deterrent. We performed the test over five consecutive days, with four days of training followed by a probe test on the fifth day. During the training period, each mouse was placed in the center of the maze in an opaque chamber for 10 s, and then allowed 3 min to explore and find the escape box. If the mouse did not find the escape box, the animal was gently guided to the target and placed inside the box for 1 min. If the mouse placed its nose in the target hole for at least 20 consecutive seconds, we recorded the mouse as entering the target box. Mice underwent four trials per training day and rested in their home cage for 15 min between trials. The latency to enter the escape box was recorded. During the probe test on the fifth day, the escape box was removed and replaced with an insert identical to the other holes of the maze. Each mouse was allowed 90 s to find the target hole. The duration spent and the number of bouts at the target hole were recorded.

### Frailty Index Assessment

The presence of aging phenotypes, or frailty index (FI), was assessed using several health-related signs of physical deterioration as previously described [16,18]. All mice were evaluated by the same researcher and were scored as follows: 0 to indicate no sign of deficit, 0.5 to rate a mild deficit, and 1 to reflect a severe deficit. The following criteria were assessed: **Integument/skin:** alopecia, loss of fur color, dermatitis, loss of whiskers, coat condition. **Physical/Musculoskeletal:** tumors, distended abdomen, kyphosis, tail stiffening, gait disorders, tremor, forelimb grip strength, body condition score. **Vestibulocochlear/Auditory:** vestibular disturbance, hearing loss. **Ocular/Nasal:** cataracts, corneal opacity, eye discharge/swelling, microphthalmia (reduction of eye volume [27]), vision loss, menace reflex (blink response to a visual threat [28]), nasal discharge. **Digestive/urogenital:** malocclusions, rectal prolapse, vaginal/uterine/penile prolapse, diarrhea. **Respiratory system:** breathing rate/depth. **Discomfort:** mouse grimace scale, piloerection. **Body surface temperature** (La Crosse Technology, La Crosse, WI, USA). **Body weight**. A frailty index score between 0 and 1 for each animal was calculated by dividing the sum by the number of criteria.

### Statistical Analysis

Data are presented as mean value (M) or percentage with SEM or confidence interval (CI) and are indicated in the figure legend together with sample size (N). Statistical analyses, nonlinear regression model, Pearson correlation, and one-way or two-way ANOVA using Tukey multiple comparison, with an α level of 0.05, were performed using appropriate software (GraphPad Prism v. 9, San Diego, CA, USA). Outliers were removed by utilizing the 90% prediction bands of the data set. Significances are denoted in figures with \**p*<0.05, \*\**p*<0.01, \*\*\**p*<0.001, and \*\*\*\**p*<0.0001, with α indicating a trend with *p*<0.10. We evaluated the data independent of sex and also whether a sex difference existed, as done previously when evaluating mouse behaviors [26].

## Results

### Frailty Index and Aging Phenotypes

Male and female C57BL/6J mice at three ages, young (3 months), middle aged (12 months), and old (24 months), were tested using a battery of behavioral and cognitive assays, including the open-field (OF), Y-maze, elevated plus maze (EPM), and Barnes maze. Subsequently, animals were evaluated using a non-invasive frailty index (FI) assessment, which provides a composite score derived from 31 clinical parameters (**Figure 1A**). Analysis revealed that FI score increased significantly in both male (**Figure 1B**) and females (**Figure 1C**) mice, as well as when data were analyzed irrespective of sex (**Figure 1D**) with advancing chronological age. These findings confirm that the FI score correlates with chronological age, consistent with previous reports [17].

**Figure 1.**
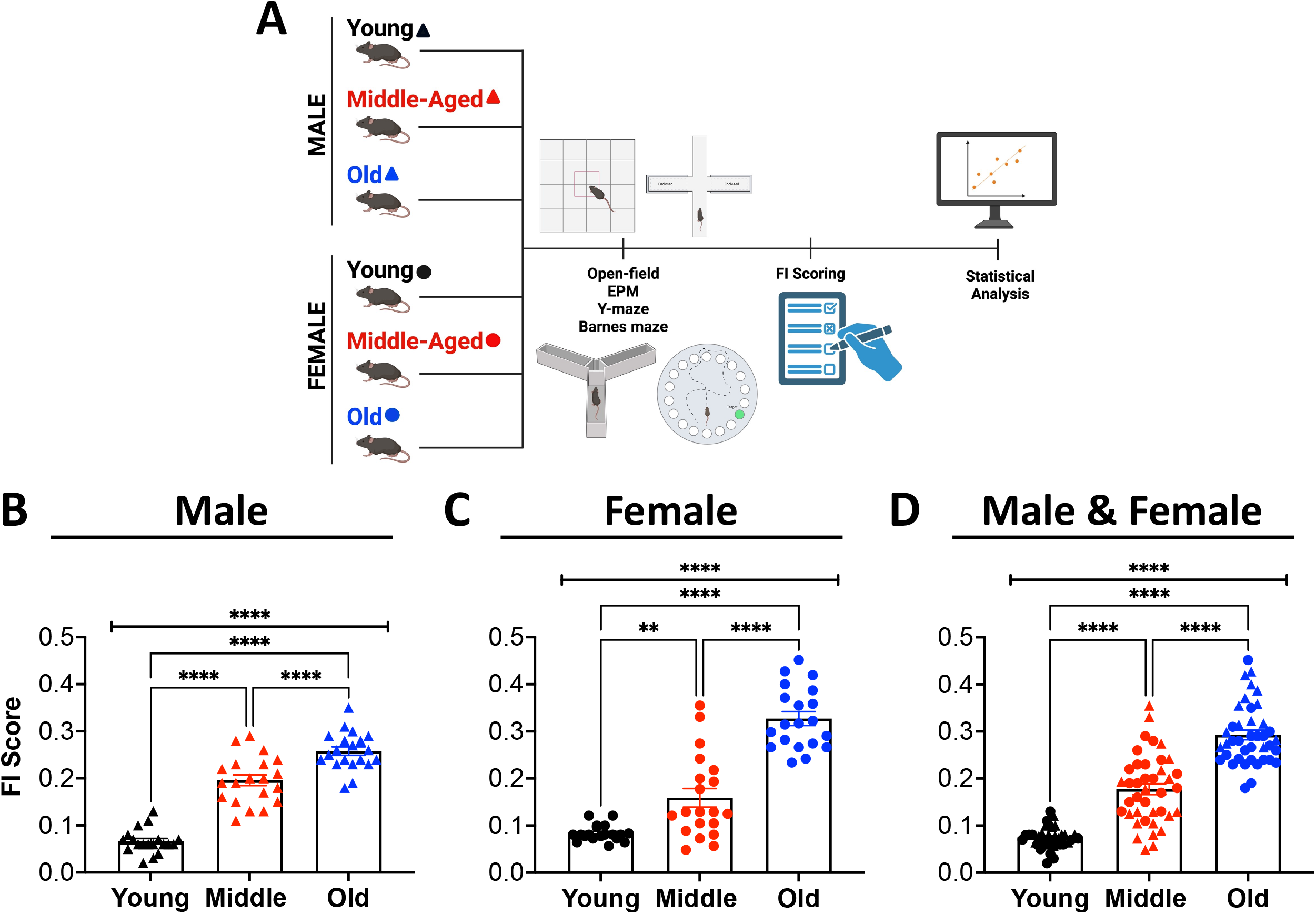
Frailty Index increases with chronological age. (A) Schematic representation of the experimental design. Mice from different age groups were assessed using the Open-Field, Elevated Plus Maze, Y-maze, and Barnes Maze behavioral assays, and overall health was evaluated using the frailty index (FI). (B–D) FI scores of C57BL/6J mice by age and sex. Male mice are represented by triangles and female mice by circles. Young (3 mos, black), middle-aged (12 mos, red), and old (24 mos, blue) mice are shown. FI scores increased significantly with age in (B) males (F = 115.1, p < 0.0001), (C) females (F = 73.1, p < 0.0001), and (D) combined sexes (F = 136.9, p < 0.0001). Sample sizes: N = 19 per sex for young, N = 20 per sex for middle-aged and old groups. Data are presented as mean ± SEM. Statistical significance was determined by one-way ANOVA with Tukey’s multiple comparisons test (**p < 0.01; ****p < 0.0001). Panel A was created in BioRender (accessed 5 September 2025).

### Frailty Index and Spontaneous Locomotion

To assess whether FI scores are associated with age-related cognitive decline, performance in behavioral assays was correlated with FI measures. Spontaneous locomotor activity and explorative behavior were examined using the open-field test, where increased percent time spent in the center of the arena and decreased total distance traveled are considered indicative of cognitive decline associated with brain aging in mice [29]. Both male and female mice exhibited increased percent time spent in the center of the open-field apparatus with advancing chronological age, an effect independent of sex (**Figure 2A–C**). Correlation analysis revealed a strong positive correlation between FI score and time spent in the center in both males (r = 0.5069) and females (r = 0.4109) (**Figure 2D–E**), as well as when data were analyzed irrespective of sex (r = 0.4106) (**Figure 2F**). These findings indicate that FI is a reliable predictor of open-field center duration and may therefore provide insight into brain aging. Analysis of total distance traveled in the OF test revealed no significant difference among age groups in either sex (**Figure S1A-C**). Correlation with FI score showed a slight negative association in males (r = −0.1556) (**Figure S1D**), and females (r = −0.1673) (**Figure S1E**), but no correlation when the two sexes were combined (r = −0.05964) (**Figure S1F**). These findings indicate that total distance over the 60-minute duration of the OF assay does not significantly change with chronological age and therefore cannot be predicted by FI scores.

**Figure 2.**
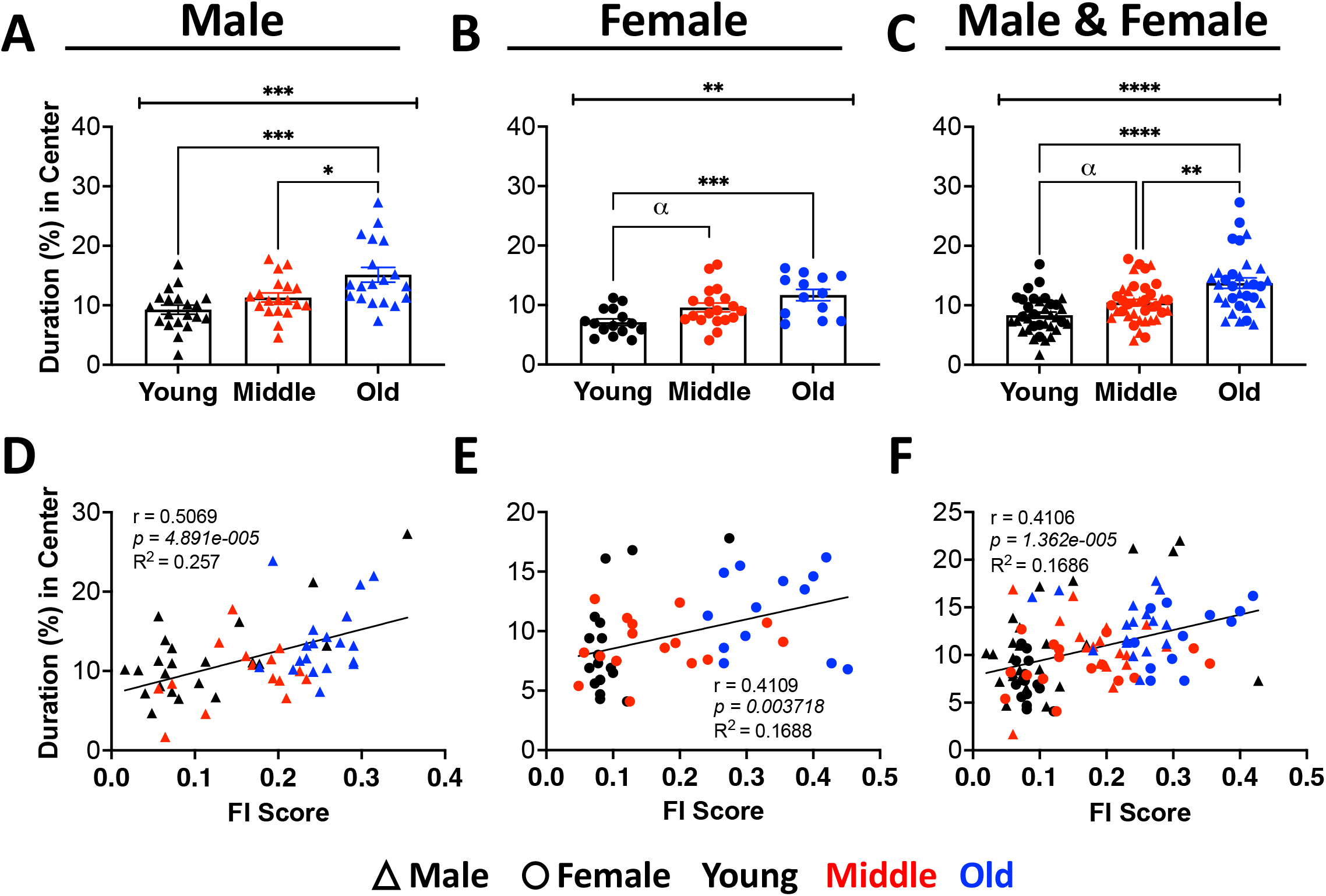
Percent duration in center of open-field test correlates with frailty index and aging. Percent duration in the center of the open-field arena and its correlation with frailty index (FI) in normal aging C57BL/6J male and female young (3 mos, black, N≥15 per sex), middle-age (12 mos, red, N≥19 per sex), and old (24 mos, blue, N≥13 per sex) mice. Percent duration in center compared to chronological age in (A) males (F=10.01, p<0.001), (B) females (F=8.083, p<0.01), as well as (C) independent of sex (F=16.63, p<0.0001). Values are percent mean ± SEM. Significances were determined using one-way ANOVA using Tukey multiple comparisons test with \**p*<0.05, \*\**p*<0.01, \*\*\**p*<0.001, and \*\*\*\**p*<0.0001. Percent duration in center correlated with FI score in (D) males (r=0.5069, p<0.0001), and (E) females (r=0.4109, p<0.0001), as well as (F) independent of sex (r=0.4106, p<0.0001). Pearson correlation coefficient (r), significance level (*p* value), and coefficient of determination (R^2^) are shown.

### Frailty Index and Anxiety-Related Assays

The elevated plus maze (EPM) is commonly employed to assess anxiety-like behavior in mice, with reduced time spent in and fewer entries into the open arms interpreted as increased anxiety-like behavior. Interestingly, aging has been associated with increased time spent in the open arms as well as a greater number of open-arm entries, a pattern that is thought to reflect reduced cognitive function rather than decreased anxiety [26][30], making it possible to study age-related changes in cognition using this test. Using this assay, we confirmed that both male and female mice spent more time in the open arms with increasing chronological age (**Figure 3A and B**), with significance reached when tested independent of sex (**Figure 3C**). Interestingly, we found a good significant positive correlation between frailty index and duration in open arms among both males (r = 0.3104) (**Figure 3D**) and females (r = 0.3033) (**Figure 3E**) as well as when both sexes were combined (r = 0.2908) (**Figure 3F**). In addition to time spent in the open arms, the total number of entries (bouts) was evaluated. We observed an age-associated increase in open-arm entries in both sexes, which reached statistical significance only in old male mice, as compared to young males (**Figure S2A–C**). Correlation analysis revealed a significant positive association between FI and the number of open-arm entries in males (r = 0.4047) and females (r = 0.3012), as well as when both sexes were combined (r = 0.3397) (**Figure S2D–F**). Further analysis within the cohort of old mice demonstrated a significant strong positive correlation between FI and both duration (r = 0.6596) and entries (r = 0.6725) in the open arms, particularly in females (**Figure S3**). These findings indicate that FI scores correlate with both the time spent and the number of entries in the open arms, reflecting age-related changes in cognitive function. Notably, old female mice exhibited a pronounced behavioral phenotype in the EPM paradigm, which shows a strong positive correlation with FI.

**Figure 3.**
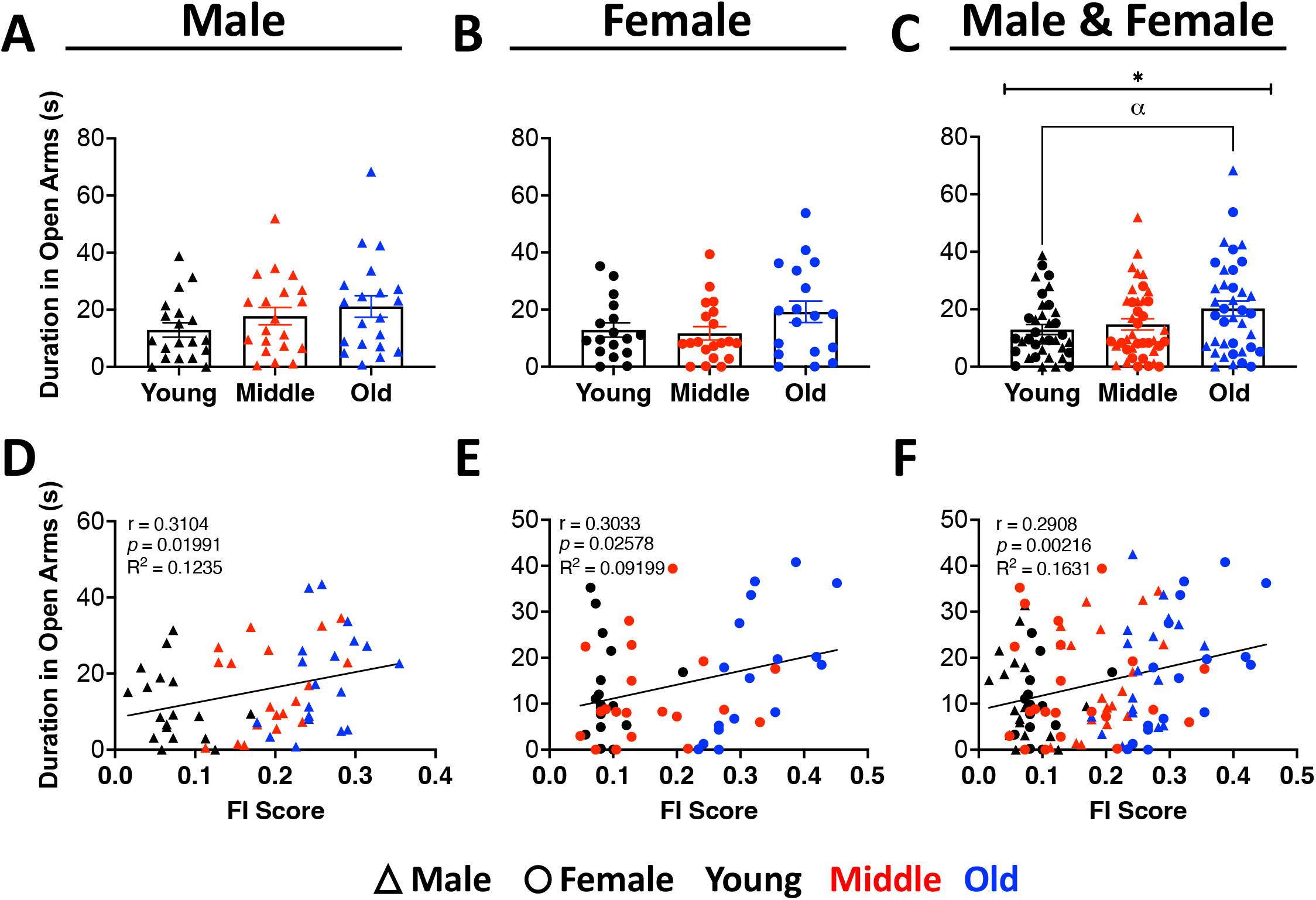
Duration in open arms of the elevated plus maze correlates with frailty index and aging. Duration in the elevated plus maze open arms and its correlation with frailty index (FI) in normal aging C57BL/6J male and female young (3 mos, black, N≥17 per sex), middle-age (12 mos, red, N=20 per sex), and old (24 mos, blue, N≥18 per sex) mice. Duration in the open arm compared to chronological age in (A) males (F=1.663, p=0.1987) and (B) females **(B)** (F=1.948, p=0.1528), as well as (C) independent of sex (F=3.086, p<0.05). Values are mean ± SEM. Significances were determined using one-way ANOVA using Tukey multiple comparisons test with \**p*<0.05, and α indicating a trend with *p*<0.10. Duration in the open arms correlated with FI score in (D) males (r=0.3104, p<0.05), and (E) females (r=0.3033, p<0.05), as well as (F) independent of sex (r=0.2908, p<0.01). Pearson correlation coefficient (r), significance level (*p* value), and coefficient of determination (R^2^) are shown.

### Frailty Index and Short-Term Spatial Learning

The Y-maze behavioral assay is commonly used to assess short-term spatial memory in mice, with reduced entries (bouts) into and time spent in the novel arm interpreted as indicators of age related cognitive decline [31]. When using this assay, we observed that entries into the novel arm decreased with advancing age in both male and female mice; however, this change reached statistical significance only in males when analyzed by sex (**Figure 4A-C**). In addition to entries, time spent in the novel arm also decreased significantly with age; however, unlike entries, this effect was significant only in female mice when analyzed by sex (**Figure S4A–C**). Correlation analysis revealed a significant negative association between FI and the number of entries into the novel arm in males (r = −0.5027) (**Figure 4D**) and when sexes were combined (r = −0.4062) (**Figure 4F**), whereas only a modest negative correlation was observed in females (r = −0.1034) (**Figure 4E**). Correlation between time spent in the novel arm and FI scores showed a moderate negative association across both sexes (male r = −0.258; female r = −0.2164) (**Figure S4D–E**) and when combined (r = −0.1587) (**Figure S4F**). These findings suggest that FI scores reliably predict age-related decline in spatial memory only in males when assessed by the number of entries into the novel arm in the Y-maze.

**Figure 4.**
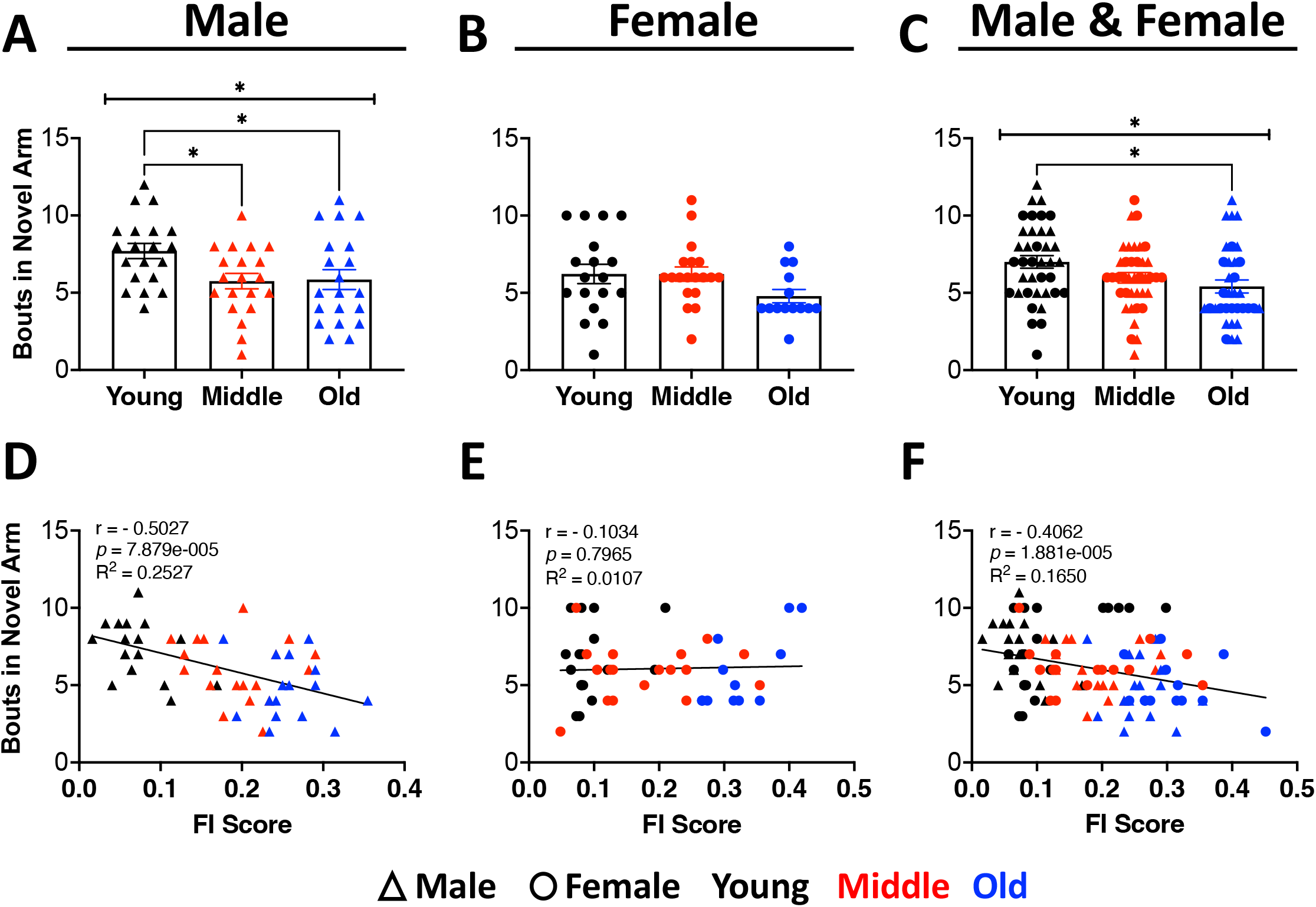
Bouts in novel arm of the Y-maze correlates with frailty index and aging. Bouts in the novel arm of the Y-maze and its correlation with frailty index (FI) in normal aging C57BL/6J male and female young (3 mos, black, N≥18 per sex), middle-age (12 mos, red, N≥19 per sex), and old (24 mos, blue, N≥14 per sex) mice. The number of bouts in the novel arm represented alongside chronological age in (A) male (F=4.014, p<0.05), (B) female (F=2.182, p=0.1239), and (C) male and female combined (F=4.224, p<0.05) groups. Values are mean ± SEM. Significances were determined using one-way ANOVA using Tukey multiple comparisons test with \**p*<0.05, and α indicating a trend with *p*<0.10. The number of bouts in the novel arm negatively correlated with FI score in (D) males (r=-0.5027, p<0.0001), (E) females (r=-0.1034, p=0.7965), as well as (F) independent of sex (r=-0.4062, p<0.0001). Pearson correlation coefficient (r), significance level (*p* value), and coefficient of determination (R^2^) are shown.

### Frailty Index and Long-Term Spatial Learning

The Barnes maze assay is commonly used to assess long-term spatial learning and memory in mice [32], with reduced time spent in the target hole and fewer entries considered indicators of cognitive decline [33]. We observed that male mice, as well as male and female mice analyzed together, spent significantly less time in the target hole and correspondingly more time in the surrounding holes with advancing age (**Figure 5A, C**), whereas female mice alone did not exhibit a significant change in target-hole duration across age groups (**Figure 5B**). Correlation analysis revealed a marked negative association between FI scores and time spent in the target hole in both males (r = −0.3578) and females (r = −0.3196) (**Figure 5D, E**), as well as when sexes were combined (r = −0.3339) (**Figure 5F**), suggesting that FI score is a good predictor of cognitive decline and brain aging assessed by target-hole duration in the Barnes maze. In addition to duration at the target hole, the number of entries into the target hole was also assessed. Here, only old male mice showed a significant reduction in target-hole entries, as compared to middle-aged males (**Figure S5A–C**). Correlation with FI scores demonstrated a significant negative association in males (r = −0.3126) (Figure S5D), a modest negative correlation in females (r = −0.2754) (**Figure S5E**), and a moderate negative correlation when sexes were combined (r = −0.2749) (**Figure S5F**). These findings indicate that duration spent in the target hole is a more sensitive measure of long-term spatial learning and memory decline with age than the number of entries.

**Figure 5.**
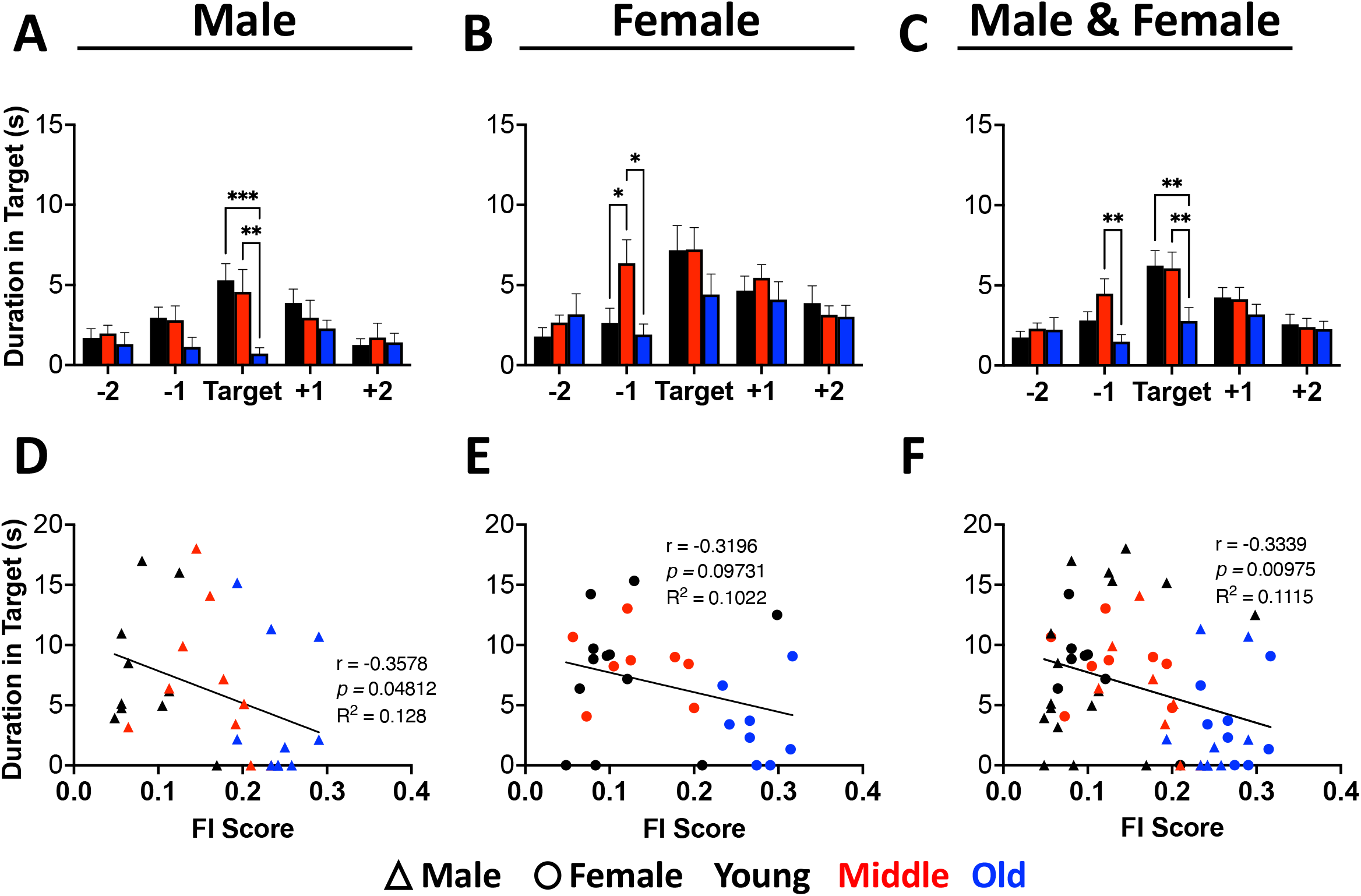
Duration in the target hole of the Barnes maze correlates with frailty index and aging. Duration in the target hole of the Barnes maze and its correlation with frailty index (FI) in normal aging C57BL/6J male and female young (3 mos, black, N≥10 per sex), middle-age (12 mos, red, N≥10 per sex), and old (24 mos, blue, N≥10 per sex) mice. The duration spent in the target hole of the Barnes maze represented alongside chronological age in (A) males (F=6.363, p<0.01), (B) females (F=6.625, p<0.01), and when evaluated (C) independent of sex (F=3.189, p<0.05). Values are mean ± SEM. Significances were determined using one-way ANOVA using Tukey multiple comparisons test with \**p*<0.05, and α indicating a trend with *p*<0.10. The duration spent in the target hole of the Barnes maze correlated with FI score in (D) males (r=-0.3578, p<0.05), (E) females (r=-0.3196, p<0.10), as well as (F) males and females combined (r=-0.3339, p<0.01). Pearson correlation coefficient (r), significance level (*p* value), and coefficient of determination (R^2^) are shown.

## Discussion

Following Horvath’s discovery in 2013, the DNA methylation clock has emerged as a reliable and valid tool for estimating biological age and predicting mortality across multiple species, including humans [10,12,34,35]. Despite its reliability, the methylation clock, along with other complex molecular assays, are typically invasive, expensive, and time consuming [11]. Consequently, many researchers have turned to the frailty index as an alternative predictor of biological aging, given its strong correlations with chronological age and diverse health outcomes [17,19]. Unlike molecular assays, the frailty index offers several advantages, such that it is easy to use, non-invasive, inexpensive, and responds to interventions [36].

Although the frailty index is widely recognized for its simplicity and utility in assessing overall aging, no studies to date have specifically investigated whether the aging phenotype scores derived from this measure can be used to predict brain aging in mice. Importantly, establishing a possible link could provide a valuable, non-invasive approach for monitoring neurodegenerative decline and brain aging. In this study, we demonstrate, for the first time, that frailty index can serve as a reliable predictor of age-related cognitive decline in mice. Cognitive performance was assessed through a series of behavioral tests, including the open-field, elevated plus maze, Y-maze, and Barnes maze behavioral paradigms, and compared against measures of physical aging such as the frailty index.

In this study, our results confirm that frailty index increases with chronological age (**Figure 1A-C**) and correlates with various behavioral outcomes commonly used in the literature to measure cognitive decline in both “normal” and pathological models of aging [33,37–40]. When investigating outcomes of the open-field behavioral paradigm, a reliable and widespread paradigm used to measure spontaneous locomotion [41], we found that as both male and female mice chronologically age and present with higher frailty index scores, they spent a larger percentage of time in the center of the open-field apparatus (**Figure 2A-F**). These data replicate and expand on previous findings that older male mice tend to spend more time in the center of the open-field arena [29]. Interestingly, we did not find a strong correlation between frailty index score and total distance traveled in the open-field area (**Figure S1A-F**). Precise motor movement is often shown to decrease with age, which may be caused by reduced strength from atrophy and a decrease in coordination [42]. On the other hand, it has been suggested that anxiety-associated behaviors can increase movement in aging mice [43]. Female mice have been traditionally shown to move more in the open-field when compared to male mice, perhaps due to the estrous cycle [44]. Moreover, the slight increase in total distance traveled by the aged female mice could be explained by an increase in anxiety-related behaviors, which tend to be two-times more prevalent in females than males [45]. Thus, these variable sex-dependent behavioral outcomes may indicate that observed changes in total distance traveled may not serve as the most reliable marker of cognitive aging.

Moreover, when investigating outcomes of the elevated plus maze (EPM) behavioral paradigm, a reliable and widespread test used to measure anxiety-like behavior, we found that as both male and female mice chronologically aged and had higher frailty index scores, these animals spent a larger percentage of time in the open arms (**Figure 3A-F**) and made fewer bouts (**Figure S2A-F**) in the EPM apparatus. Interestingly, when comparing only old mice of both sexes we identified an even stronger correlation between frailty index score and duration in the open arms of the EPM (**Figure S3A-F**). These data replicate previous findings that older mice tend to spend a larger percentage of time and have a larger percentage of bouts in open-arms of the EPM [26]. Our findings demonstrate that the association between frailty index and EPM parameters are more strongly correlated in male mice compared to females, and replicates previous findings that females tend to exhibit less anxiety-like behaviors in the EPM behavioral assay when compared to males [46,47].

Lastly, to investigate frailty index scores with spatial learning and memory measures, our study employed the Y-maze and Barnes maze behavioral paradigms. In the Y-maze, we found that this assay is a good indicator of age-related cognitive decline in male but not in female mice, as indicated by fewer bouts in the males (**Figure 4A**) and less time spent in the novel arm of the apparatus (**Figure S4A**). Interestingly, we found a strong negative correlation between the number of entries in the novel arm and FI in male mice (**Figure 4D**), suggesting that FI is a good predictor of male mice performance in this test during aging. Lastly, using the Barnes Maze, we found that as both male and female mice chronologically age and present with higher frailty index scores these animals spent significantly less time (**Figure 5A-F**) and exhibited fewer bouts (**Figure S5A-F**) in the target hole of the Barnes maze on probe day (when the escape box is removed). Importantly these findings align with previous data illustrating that mice exhibit significant impairments in spatial learning and memory with aging as seen in the Y-maze and Barnes maze paradigms [48,49].

Importantly, integrating frailty index measurements with the behavioral assays used in this study did not result in any adverse health effects in the animals, emphasizing the safety and practicality of this approach. It is important to note that the frailty index assessments must be consistently performed by the same researcher to ensure reliability. Additionally, there are various frailty index assessments available and utilized by groups to investigate frailty in mice; therefore, a different FI scoring system may yield different findings and it is important for researchers to choose the correct assessment for the study at hand [50].

Taken together, our findings demonstrate that the frailty index is a valuable and non-invasive tool for exploring the link between brain aging and overall physiological aging. When combined with other aging biomarkers, such as DNA methylation profiles, this method may be beneficial for developing accurate predictions of how chronological age affects normal physiological processes, including cognitive functioning.

## Acknowledgements

We are grateful for the assistance of the animal care staff at the Harvard Medical School. Figure 1A was created using BioRender.

## Author Contributions

This manuscript is included in the doctoral thesis of Sydney Bartman. G.C. and J.M.R. conceived the study and designed the analyses. A.K., E.P., and G.C., performed experiments and S.B., L.G., G.C., and J.M.R. analyzed the data. D.A.S. and J.M.R. provided funding. S.B., L.G., G.C., and J.M.R. wrote the manuscript. E.P. contributed to preliminary data analysis and early drafts of the manuscript. All authors have read and approved the final manuscript.

## Institutional Review Board Statement

The animal study protocol was approved by the Institutional Review Board of Harvard Medical School under D.A.S..

## Informed Consent Statement

Not applicable.

## Data Availability Statement

Data generated from this study are available upon request.

## Conflicts of Interest

The authors declare that the research was conducted in the absence of any commercial or financial relationships that could be construed as a potential conflict of interest. The funders had no role in the design of the study; in the collection, analyses, or interpretation of data; in the writing of the manuscript; or in the decision to publish the results. For conflicts related to D.A.S., including his ownership and board membership of Life Biosciences, an epigenetic reprogramming company, see: https://genetics.med.harvard.edu/sinclairtest/people/sinclair-other.php

## Funding Statement

This work was supported by the National Institute on Aging (K99/R00AG055683 to J.M.R.), the Roddy Foundation (G.C. and J.M.R), the George & Anne Ryan Institute for Neuroscience (G.C. and J.M.R), the College of Pharmacy at the University of Rhode Island (G.C., J.M.R.), the Interdisciplinary Neuroscience Program (S.B.) at the University of Rhode Island, and the University of Rhode Island Dean’s Fellowship (S.B.). G.C. and J.M.R. are supported by the National Institutes of Health Office of the Director (R21OD037651). A.E.K. is supported by National Institutes on Aging (K99/R00AG070102). D.A.S is supported by the National Institute on Aging (R01AG082737), the Paul F. Glenn Foundation, the Aoki Foundation, the Centurion Foundation, the Rosenkranz Foundation, and Friends of Sinclair Lab.

