## Supplementary Material for "Associations Between Frailty and Cognitive Outcomes Across the Lifespan of Mice"

### Supplementary Figures

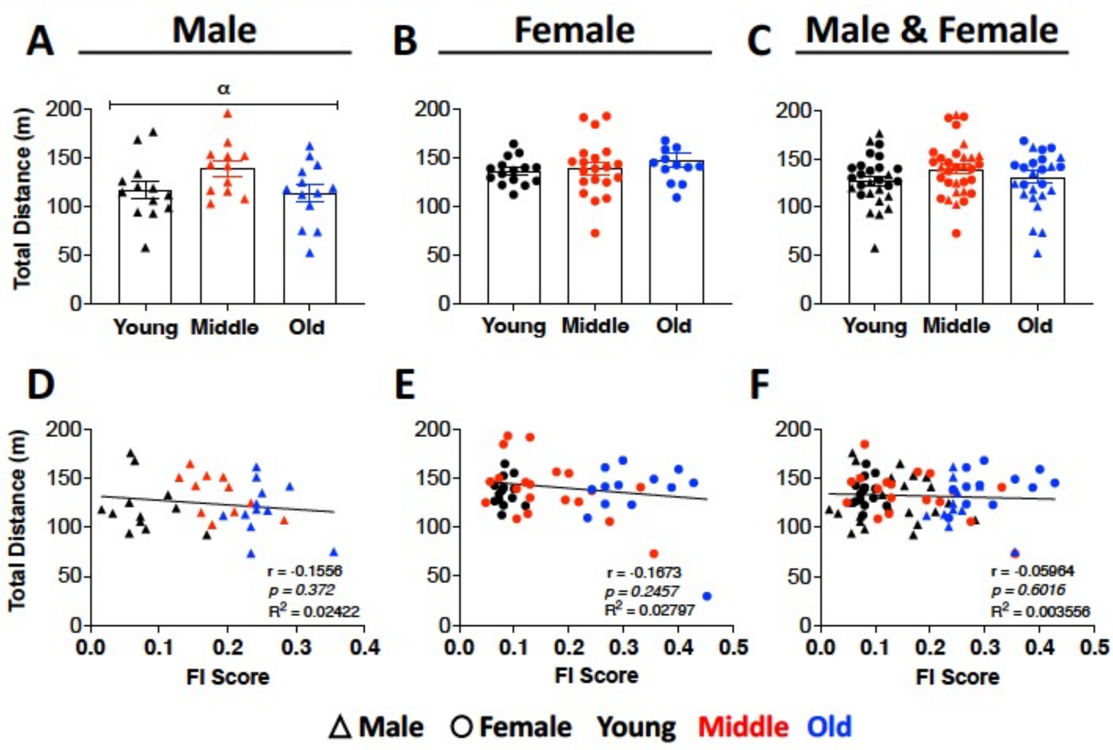


**Figure S1. Total distance in the open-field paradigm does not correlate with frailty index during aging.** Total horizontal movements over the 60-minute duration of the open-field assay and its correlation with frailty index (FI) in normal aging C57BL/6J male and female young (3 mos, black, N≥13 per sex), middle-age (12 mos, red, N≥12 per sex), and old (24 mos, blue, N≥13 per sex) mice. Total distance traveled compared with chronological age in (A) male (F=2.578, p<0.10), (B) female (F=0.7729, p=0.4677), and (C) male and female combined (F=1.408, p=0.2503) groups. Values are mean ± SEM. Significances were determined using one-way ANOVA using Tukey multiple comparisons test. Total distance correlated with FI score in (D) males (r=-0.1556, p=0.372), (E) females (r=-0.1673, p=0.2457), and with (F) both sexes combined (r=-0.05964, p=0.6016). Pearson correlation coefficient (r), significance level (*p* value), and coefficient of determination ($R^2$) are shown.

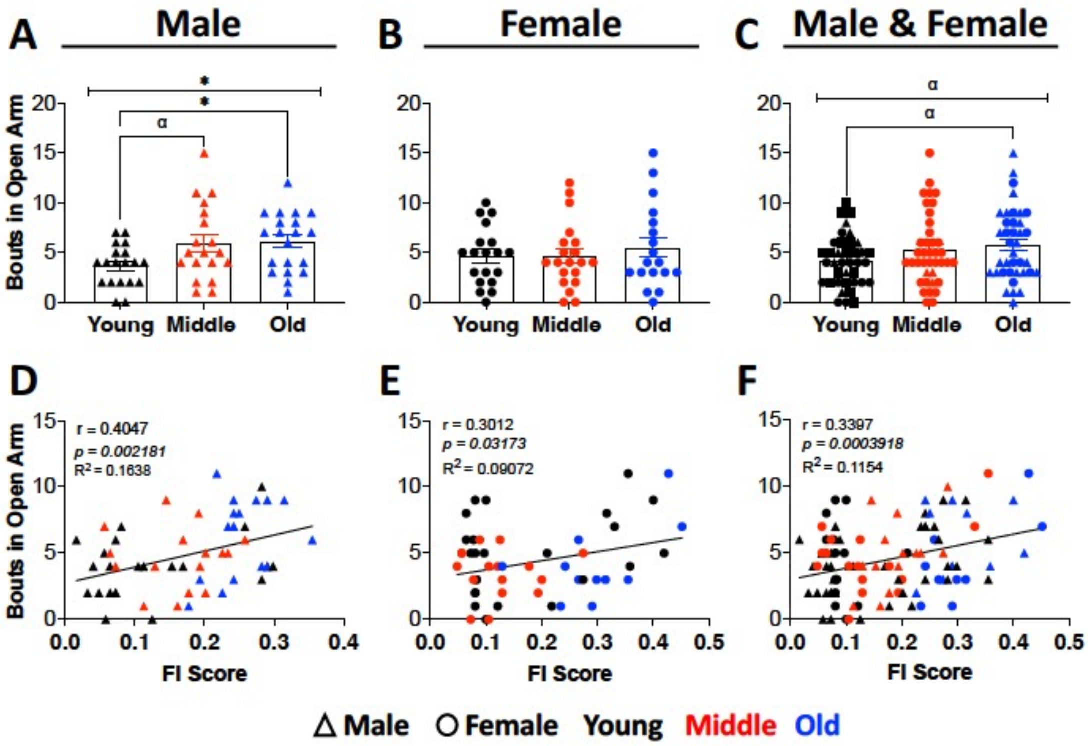


**Figure S2. Bouts in the open arms of the elevated plus maze loosely correlate with frailty index during aging.** Bouts in the open arms of the elevated plus maze and its correlation with frailty index (FI) in normal aging C57BL/6J male and female young (3 mos, black, N=19 per sex), middle-age (12 mos, red, N≥19 per sex), and old (24 mos, blue, N≥18 per sex) mice. The number of bouts in the open arms compared to chronological age in **(A)** male (F=6.086, p<0.05), (B) female (F=0.3677, p=0.6940), and (C) male and female combined (F=2.545, p<0.10) mice. Values are mean ± SEM. Significances were determined using one-way ANOVA using Tukey multiple comparisons test with *$p$<0.05, and α indicating a trend with $p$<0.10. The number of bouts in the open arms correlated with FI score in (D) males (r=0.4047, p<0.01), (E) females (r=0.3012, p<0.05), as well as (F) independent of sex (r=0.3397, p<0.001). Pearson correlation coefficient (r), significance level ($p$ value), and coefficient of determination ($R^2$) are shown.

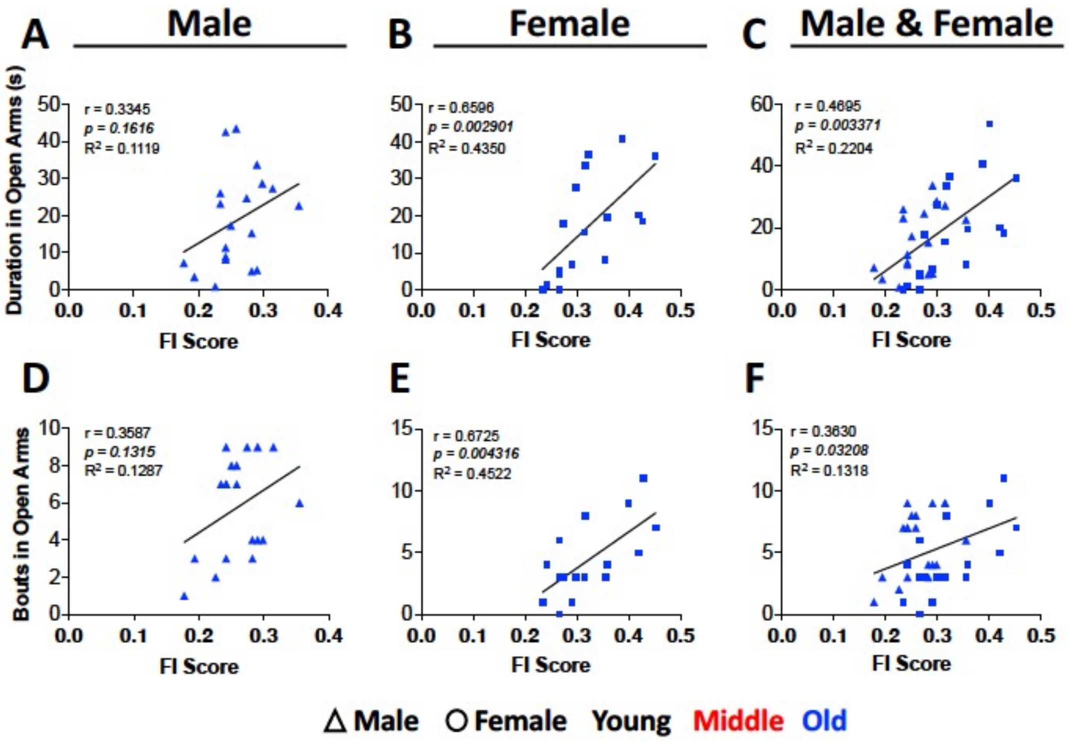


**Figure S3. Duration and number of bouts in open arms of the elevated plus maze strongly correlates with frailty index in old mice.** Duration and number of bouts in the elevated plus maze open arms and its correlation with frailty index (FI) in normal aging C57BL/6J male and female old (24 mos, blue, N≥16 per sex) mice. Duration in the open arms correlated with FI score in (A) male (r=0.3345, p=0.1616), (B) females (r=0.6596, p<0.01), as well as (C) independent of sex (r=0.4695, p<0.01). The number of bouts in the open arms correlated with FI score in (D) male (r=0.3587, p=0.1315), (E) female (r=0.6725, p<0.01) and (F) male and female combined (r=0.3630, p<0.05), mice. Pearson correlation coefficient (r), significance level (*p* value), and coefficient of determination ($R^2$) are shown.

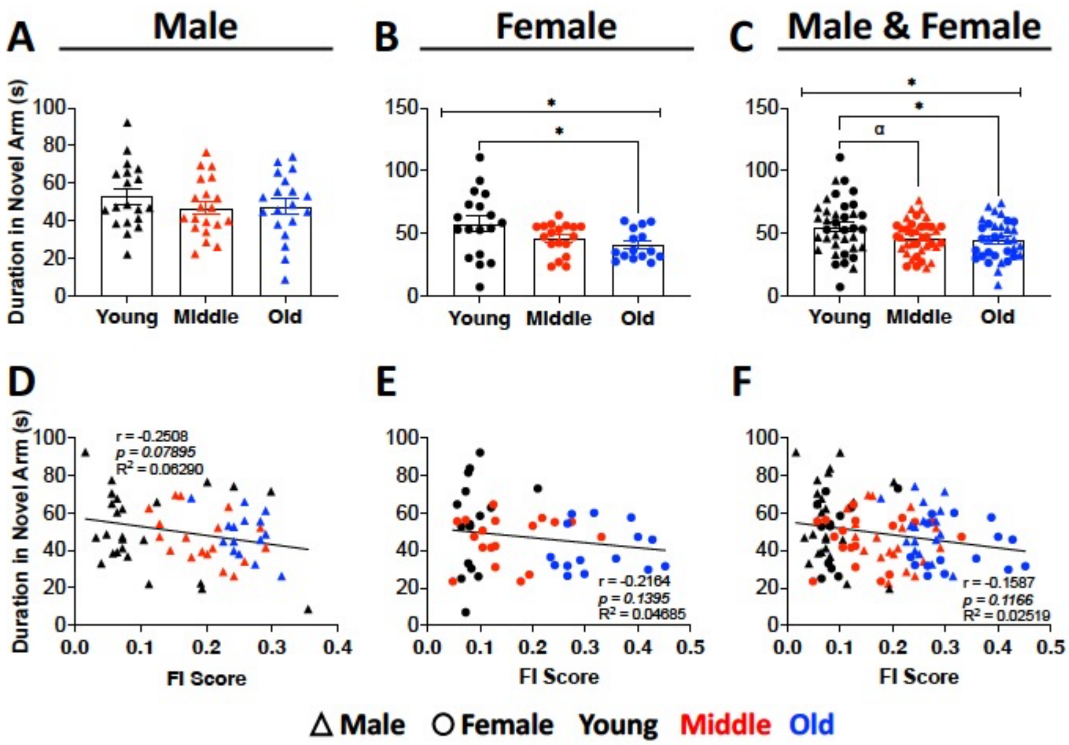


**Figure S4. Duration in novel arm of the Y-maze loosely correlates with frailty index and aging.** Duration spent in the novel arm of the Y-maze and its correlation with frailty index (FI) in normal aging C57BL/6J male and female young (3 mos, black, N≥18 per sex), middle-age (12 mos, red, N≥18 per sex), and old (24 mos, blue, N≥15 per sex) mice. The duration spent in the novel arm with chronological age in (A) males (F=0.7217, p=0.4905), and (B) females (F=3.550, p<0.05), as well as (C) males and females combined (F=3.687, p<0.05). Values are mean ± SEM. Significances were determined using one-way ANOVA using Tukey multiple comparisons test with **p*<0.05, and α indicating a trend with *p*<0.10. The duration spent in the novel arm correlated with FI score in (D) male (r=-0.2508, p<0.10), (E) female (r=-0.2164, p<0.1395), and (F) male and female combined (r=-0.1587, p=0.1166) mice. Pearson correlation coefficient (r), significance level (*p* value), and coefficient of determination ($R^2$) are shown.

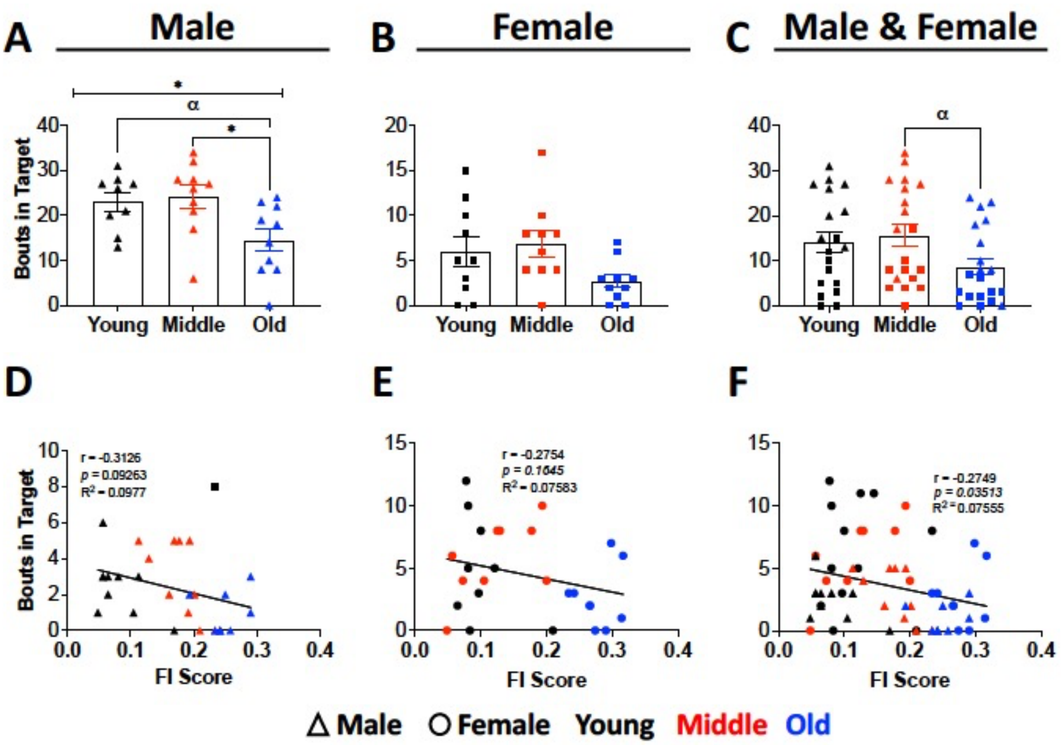


**Figure S5. Bouts in the target hole of the Barnes maze weakly correlates with frailty index and aging.** The number of bouts in the target hole of the Barnes maze and its correlation with frailty index (FI) in normal aging C57BL/6J male and female young (3 mos, black, N≥9 per sex), middle-age (12 mos, red, N≥10 per sex), and old (24 mos, blue, N≥10 per sex) mice. The number of bouts in the target hole of the Barnes maze compared to chronological age in (A) male (F=4.848, p<0.05), (B) female (F=2.791, p<0.10), and (C) male and female combined (F=2.673, p<0.10) mice. Values are mean ± SEM. Significances were determined using one-way ANOVA using Tukey multiple comparisons test with *$p$<0.05, and α indicating a trend with $p$<0.10. The number of bouts in the target hole of the Barnes maze correlated with FI score in (D) males (r=-0.3126, p<0.10), (E) females (r=-0.2754, p=0.1645), as well as (F) independent of sex (r=-0.2749, p<0.05). Pearson correlation coefficient (r), significance level ($p$ value), and coefficient of determination ($R^2$) are shown.
